# Social stress induces autoimmune responses against the brain to promote stress susceptibility

**DOI:** 10.1101/2022.11.18.517081

**Authors:** Yusuke Shimo, Flurin Cathomas, Hsiao-yun Lin, Kenny L Chan, Lyonna F. Parise, Long Li, Carmen Ferrer-Pérez, Sara Costi, James W. Murrough, Scott J Russo

## Abstract

Clinical studies have revealed a high comorbidity between autoimmune and psychiatric disorders, including major depressive disorder (MDD). However, the mechanisms connecting autoimmunity and depression remain unclear. Here, we aim to identify the processes linking adaptive immune abnormalities and depression. To examine this relationship, we analyzed antibody responses and autoimmunity in the chronic social defeat stress (CSDS) model in mice, and in clinical samples from patients with MDD. We show that socially stressed mice have elevated serum antibody concentrations. Activation of social stress-induced antibody responses were confirmed by detecting expansion of specific T and B cell populations particularly in the cervical lymph nodes, where brain-derived antigens are preferentially delivered. IgG antibody concentrations in the brain were significantly higher in stress-susceptible mice than in unstressed mice, and positively correlated with social avoidance. IgG antibodies accumulated around the blood vessels in brain sections from stress-susceptible mice. Moreover, sera from stress-susceptible mice exhibited high reactivity against brain tissue, and brain-reactive IgG antibody levels positively correlated with depression-like behavior. Similarly, in humans, increased peripheral levels of brain-reactive IgG antibodies were associated with increased anhedonia. Furthermore, high stress-resilience was observed in B cell-depleted mice, confirming a causal link between antibody-producing cells and depression-like behavior. This study provides novel mechanistic insights connecting stress-induced autoimmune reactions against the brain and stress susceptibility. Therapeutic strategies targeting autoimmune responses can therefore be devised to treat patients with MDD featuring immune abnormalities.

## Introduction

Major depressive disorder (MDD) affects 6% of adults worldwide every year (1, 2). Despite the availability of effective antidepressants and psychotherapies, more than one-third of patients with MDD are resistant to these treatments (3, 4). Such poor treatment outcomes can be ascribed to the heterogeneity of patients with MDD and lack of knowledge about causal mechanisms responsible for MDD symptoms. Recent reports have revealed that immune abnormalities can be detected in subpopulations of patients with MDD (5–9). Under physiological conditions, the immune system protects against infection and eliminates foreign substances via sequential and coordinated responses called innate and adaptive immunity (10). Innate immune responses are mediated by leukocyte populations, such as monocytes, granulocytes, and dendritic cells, which rapidly and nonspecifically react to pathogens, and eliminate them via several mechanisms, including induction of inflammation. Adaptive immune responses, mediated by two major populations of lymphocytes, T and B cells, react in a slow but specific manner. One of the most important functions of the adaptive immune responses is the production of antigen-specific antibodies from B cells.

Stress, which is a major risk factor for depression, induces activation of innate immune responses (11). Innate immune abnormalities have been associated with depression (12), and contributions of innate immune cells such as monocytes to behavioral deficits have been reported in mouse stress models (13–17). Chronic social defeat stress (CSDS) in mice induces behavioral abnormalities that partly resemble clinical symptoms of depression. In the CSDS model, after repeated exposures to social defeat stress, susceptible (SUS) mice show social avoidance, whereas resilient (RES) mice are devoid of such behavior (18). We have observed elevated pro-inflammatory cytokine IL-6 levels and blood–brain barrier (BBB) dysfunction in the CSDS model and patients with MDD (19, 20). Although several reports suggest the involvement of the adaptive immune system in neurobehavioral disorders (21–23), the specific contributions of adaptive immune abnormalities to depression remain unclear.

Clinical studies have shown a high comorbidity between psychiatric and autoimmune disorders (24). Autoimmune reactions against the brain have been implicated in the pathogenesis of psychiatric symptoms such as psychosis and cognitive impairment (25, 26). Mechanisms of autoantibody-induced neurobehavioral abnormalities are gradually becoming elucidated via the analyses of several autoimmune diseases that display psychiatric symptoms and autoantibodies against proteins expressed in the brain (27–30). However, the role of adaptive immune system dysfunction, including autoimmunity, in the pathogenesis of depression remains unclear. To understand the link between adaptive immune abnormalities and depression, we analyzed adaptive immune processes in the CSDS model and clinical samples from patients with MDD. We first investigated whether or not antibody production, which is one of the major functions of the adaptive immune system, is enhanced by stress. We further examined effects of social stress on immune cell populations controlling antibody production in lymphoid organs such as lymph nodes and the spleen (SPL).

Autoantibodies are deposited in affected organs of patients with several autoimmune diseases (31). In addition, the presence of autoantibodies targeting antigens expressed in the brain in cerebrospinal fluid (CSF) or serum is a clinical sign of some neurological diseases (25). Therefore, we explored whether antibody levels in the brain are increased in stressed mice, and if brain-reactive antibodies are found in sera from stressed mice and patients with MDD. B cell depletion therapies, such as anti-CD20 antibodies, rituximab, ocrelizumab, and ofatumumab, have been tested in clinical trials or approved for the treatment of some autoimmune diseases (32). Thus, we investigated the causal link between antibody-producing cells and stress susceptibility by depleting B cells in the CSDS model.

In this study, by analyzing adaptive immune responses in the CSDS model, we found elevation of serum antibody levels and strong induction of antibody responses in the brain-draining lymph nodes from SUS mice. Brain-reactive antibodies were induced by stress and correlated with depression-like behavior, suggesting that autoimmune responses against the brain contribute to stress susceptibility. We observed a similar association between high brain-reactivity of sera and anhedonia in humans, which may indicate clinical relevance of the findings from the CSDS model.

## Results

### Social Stress Induces Antibody Production

To explore potential dysfunction of adaptive immune responses in a mouse model of chronic psychosocial stress, C57BL/6J mice underwent a 10-day social defeat period followed by social interaction (SI) testing (**Fig. 1A**). Stressed mice were classified as SUS or RES based on their SI ratio (**Figs. 1B and C**). Although both SUS and RES mice moved shorter distances during the SI test in the absence of a social target when compared with unstressed control (CON) mice, no significant differences were observed in locomotion between SUS and RES mice (**Fig. 1D**). We then collected peripheral blood samples 48 h after the last bout of defeat and analyzed total IgG antibody concentrations in sera via an enzyme-linked immunosorbent assay (ELISA). The sera from SUS mice showed significantly higher IgG antibody concentrations than the sera from CON mice (**Fig. 1E**). In addition, IgG concentrations negatively correlated with the SI ratio (**Fig. 1F**). These results suggest that social stress induces antibody production, which may contribute to social avoidance behavior.

**Fig. 1.**
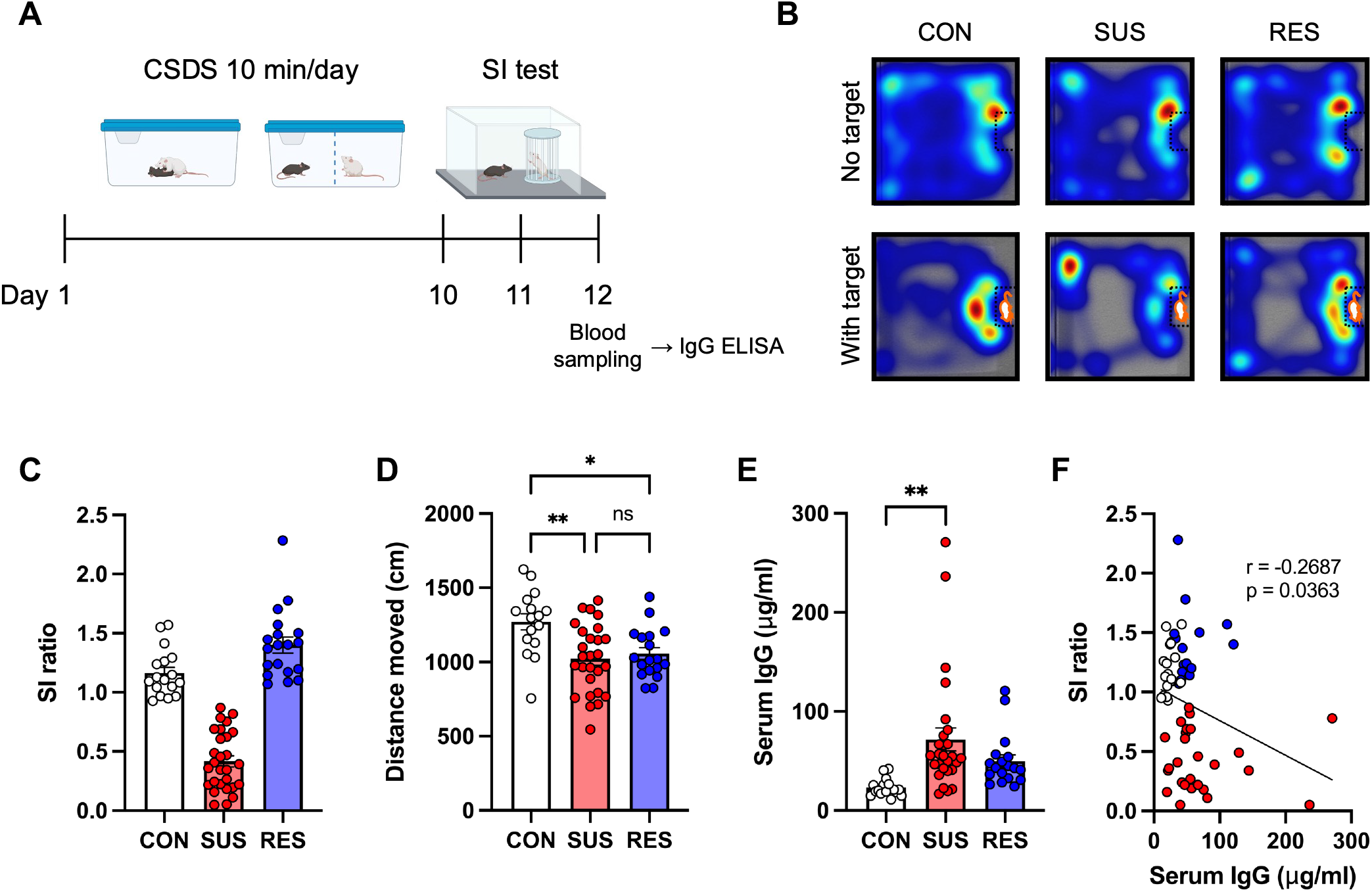
Increased antibody production in stress-susceptible (SUS) mice. **A** Experimental outline: 10 days of chronic social defeat stress (CSDS) followed by social interaction (SI) test, blood sampling, and enzyme-linked immunosorbent assay (ELISA). **B** Representative heatmaps of the unstressed control (CON), SUS, and stress-resilient (RES) mouse behavior during the SI test. **C** Social behavior in the CSDS model assessed by the SI test (CON: n = 16, SUS: n = 27, RES: n = 18). **D** Distance moved during the SI test (No target) (CON: n = 16, SUS: n = 27, RES: n = 18). **E** Total IgG antibody concentrations in sera (CON: n = 16, SUS: n = 27, RES: n = 18). **F** Correlation between SI ratio and serum IgG antibody concentrations (CON: n = 16, SUS: n = 27, RES: n = 18). Data represent mean ± standard error of the mean. One-way ANOVA with Bonferroni *post hoc* test (*p < 0.05, **p < 0.01, ns: not significant). Correlation was evaluated by Pearson correlation analysis.

### Social Stress Induces Antibody Responses Preferentially in Brain-draining Lymph Nodes

T cell-dependent antibody responses are primarily induced in organized structures in secondary lymphoid organs called germinal centers. Follicular helper T cells (Tfh), which promote B cell differentiation and proliferation, germinal center B cells (GCB), and plasma cells (PC), the actual antibody-producing cells, are involved in the germinal center reaction (33). Tissue-derived antigens are preferentially delivered to specific lymphoid organs, such as lymph nodes and the SPL. Recent reports suggest the involvement of the gut–brain axis in depression (34, 35). Gut-associated lymphoid tissues, such as mesenteric lymph nodes (mLN) and Peyer’s patches, are sites where gut immune responses are induced. The SPL captures blood-derived antigens, and splenic immune cells are activated in mouse models which induce depression-like behaviors (36, 37). Cervical lymph nodes (cLN) have been identified as sites of antigen delivery from the brain via lymphatic vessels in the meninges (38, 39). To investigate whether antibody responses were induced in specific lymphoid organs after CSDS, we analyzed immune cell populations involved in the germinal center reaction. cLN, mLN, and SPL were collected 48 h after the last defeat, and analyzed for GCB, Tfh, and PC populations via flow cytometry (**Figs. 2A, B and S1**). We observed a significant increase in the percentage of GCB only in the cLN from SUS mice (**Figs. 2C–E**). Following CSDS, the percentages of Tfh significantly increased in the cLN and mLN but not in the SPL. Notably, cLN were the only lymphoid organs which showed a significant increase in Tfh only in SUS mice compared with that in CON mice (**Figs. 2F–H**). In particular, the percentage of Tfh in the cLN from SUS mice was significantly higher than that in both RES and CON mice (**Fig. 2F**). Meanwhile, the percentage of PC increased in all the analyzed lymphoid organs, particularly in the cLN from SUS mice, showing a 17-fold increase compared with that in the cLN from CON mice (**Figs. 2I–K**). In addition, correlations between the immune cell population frequencies and depression-like behavior were the most pronounced in the cLN (**Figs. 2L–N and S2A–F**). Together, these results highlight that social stress strongly induces antibody responses in brain-draining lymph nodes.

**Fig. 2.**
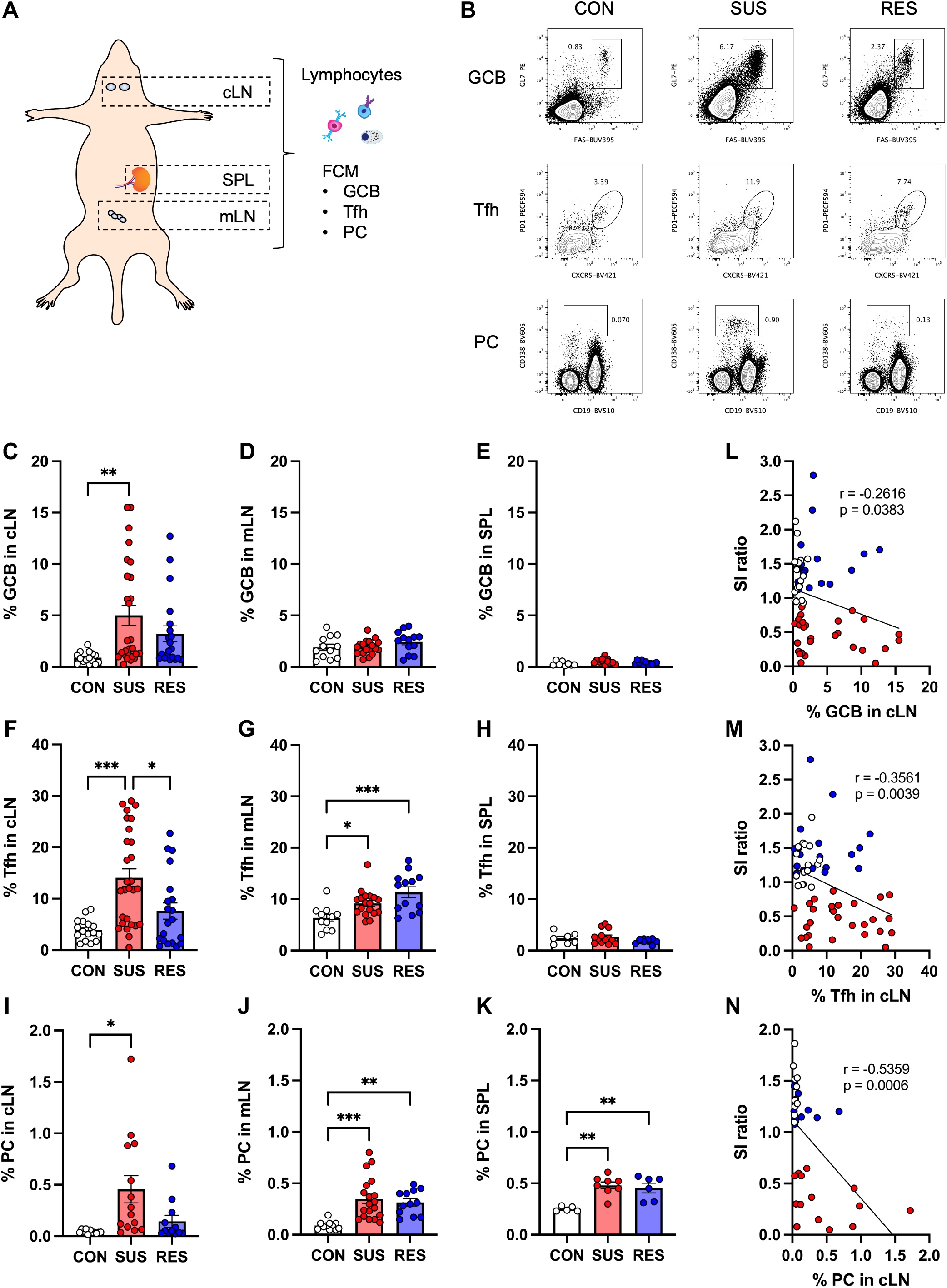
Preferential induction of immune cell populations controlling the germinal center reaction in brain-draining lymph nodes from stress-susceptible (SUS) mice. **A** Anatomical diagram showing the locations of lymphoid organs assessed by flow cytometry (FCM). **B** Representative flow cytometry plots of germinal center B cells (GCB), follicular helper T cells (Tfh), and plasma cells (PC) in cervical lymph nodes (cLN) from unstressed control (CON), SUS, and stress-resilient (RES) mice. (C-K) Percentages of GCB in **C** cLN (CON: n = 15, SUS: n = 27, RES: n = 20), **D** mesenteric lymph nodes (mLN) (CON: n = 12, SUS: n = 18, RES: n = 13), and **E** spleen (SPL) (CON: n = 8, SUS: n = 12, RES: n = 11). Percentages of Tfh in **F** cLN (CON: n = 16, SUS: n = 28, RES: n = 20), **G** mLN (CON: n = 11, SUS: n = 18, RES: n = 13), and **H** SPL (CON: n = 7, SUS: n = 13, RES: n = 10). Percentages of PC in **I** cLN (CON: n = 11, SUS: n = 14, RES: n = 12), **J** mLN (CON: n = 11, SUS: n = 19, RES: n = 12), and **K** SPL (CON: n = 5, SUS: n = 8, RES: n = 6). (L-N) Correlation between social interaction (SI) ratio and percentages of **L** GCB (CON: n = 16, SUS: n = 27, RES: n = 20), **M** Tfh (CON: n = 16, SUS: n = 28, RES: n = 20), and **N** PC (CON: n = 11, SUS: n = 14, RES: n = 12) in cLN. Data represent mean ± standard error of the mean. One-way ANOVA with Bonferroni *post hoc* test (*p < 0.05, **p < 0.01, ***p < 0.001) Correlations were evaluated by Pearson correlation analysis.

### IgG Antibodies Accumulate in the Brain Vasculature after CSDS

Considering the preferential expansion of the T and B cell populations in brain-draining lymph nodes, we hypothesized that social stress induces antibody responses against antigens expressed in the brain, consequently promoting stress susceptibility. To test this, we prepared brain lysates and sections from PBS-perfused mice (to remove potential IgG antibodies from circulation) after CSDS and analyzed IgG antibodies in the brain quantitatively by ELISA and qualitatively using immunohistochemistry (IHC) (**Figs. 3A, S3A, and B**). Brain lysates from SUS mice showed significantly higher IgG antibody concentrations than those from CON mice (**Fig. 3B**). Furthermore, the IgG antibody concentrations in the brain negatively correlated with social behavior (**Fig. 3C**). Interestingly, the levels of IgG antibodies in the brain clearly correlated with the percentages of immune cell populations controlling antibody production in the cLN (**Figs. 3D–F**). Next, brain sections containing the nucleus accumbens (NAc) (**Fig. 3G**), hippocampus (HIP), and prefrontal cortex (PFC) (**Fig. S3C**) from each group were stained with anti-mouse IgG secondary antibodies to detect antibodies bound to the brain. To visualize antibody localization, we co-stained these sections for components of the neurovascular unit, which consists of neurons, vascular cells (endothelial cells, pericytes, and vascular smooth muscle cells), and glial cells (astrocytes, microglia, and oligodendrocytes) (40). 3D reconstruction images from the NAc showed IgG antibodies were primarily detected in areas co-stained with cells of the vascular system, such as CD31-positive endothelial cells and GFAP-positive astrocytes in brain sections from SUS mice (**Figs. 3H and I**). These data suggest that CSDS induces brain-reactive antibodies which may contribute to the stress susceptibility.

**Fig. 3.**
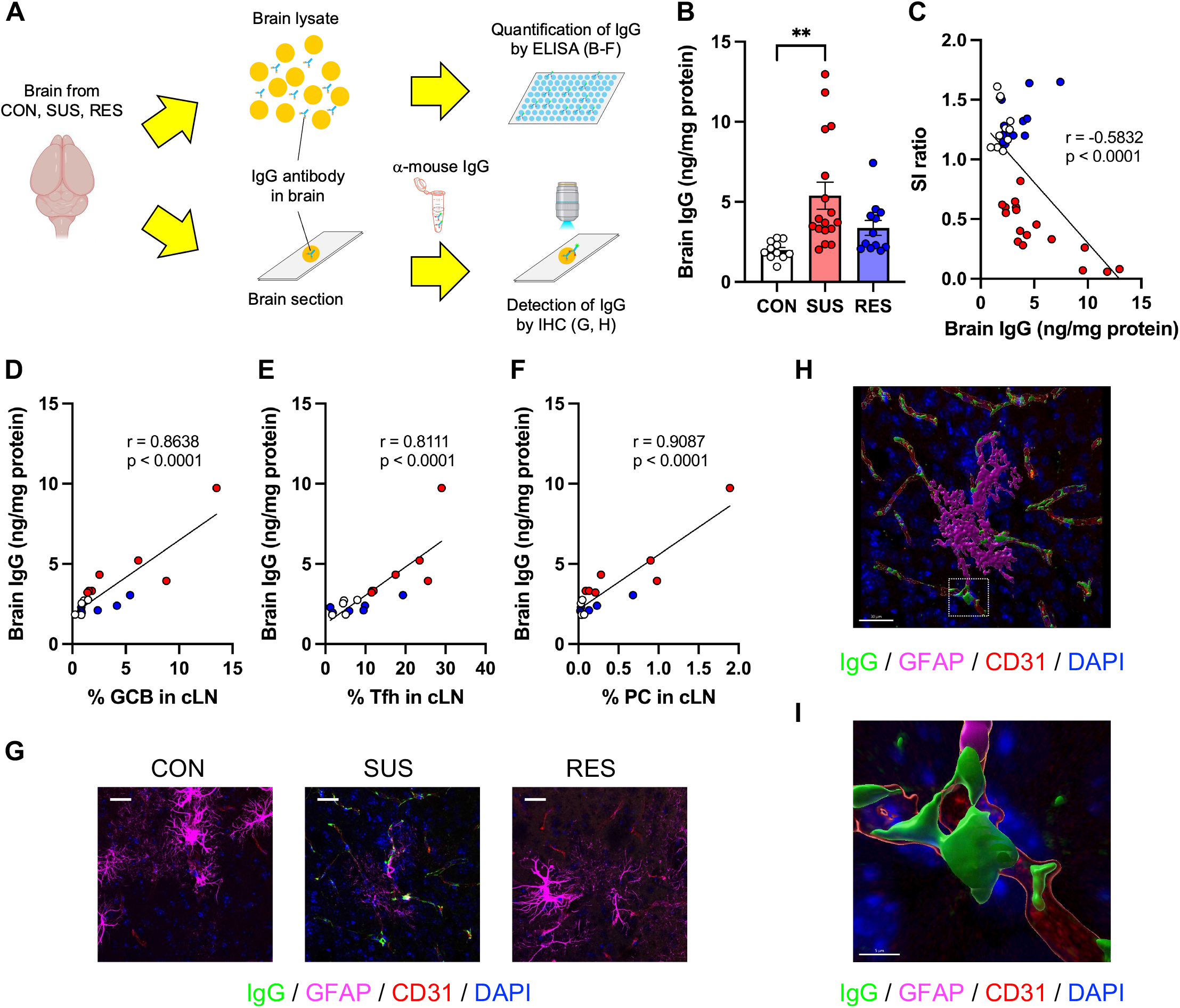
Accumulation of IgG antibodies in the brain from stress-susceptible (SUS) mice. **A** Experimental outline: Schematic of the analysis of IgG antibodies in the brain by enzyme-linked immunosorbent assay (ELISA) and immunohistochemistry (IHC). **B** Quantification of IgG antibody concentrations in brain lysates from unstressed control (CON), SUS, and stress-resilient (RES) mice (CON: n = 11, SUS: n = 17, RES: n = 12). **C** Correlation between social interaction (SI) ratio and brain IgG antibody concentrations (CON: n = 11, SUS: n = 17, RES: n = 12). (D-F) Correlations between brain IgG antibody concentrations and percentages of **D** germinal center B cells (GCB), **E** follicular helper T cells (Tfh), and **F** plasma cells (PC) in cervical lymph nodes (cLN) (CON: n = 5, SUS: n = 7, RES: n = 6). **G** Detection of IgG antibodies in brain sections around nucleus accumbens (NAc) from CON, SUS, and RES mice (green: IgG, magenta: GFAP, red: CD31, blue: DAPI, scale bar: 25 μm). (H and I) 3D reconstruction images of a brain section form a SUS mouse showing the IgG staining colocalized with brain vascular cell markers (green: IgG, magenta: GFAP, red: CD31, blue: DAPI, scale bar: **H** 30 μm, **I** 5 μm). The area inside the white frame in H is shown in I. Data represent mean ± standard error of the mean. One-way ANOVA with Bonferroni *post hoc* test (**p < 0.01). Correlations were evaluated by Pearson correlation analysis.

### Brain-reactive Antibodies Are Induced in Sera from SUS Mice and Humans with High Anhedonia

To investigate the presence of brain-reactive antibodies in sera, we quantified brain-reactive antibodies in the sera from the CSDS model using ELISA (**Figs. 4A, S4A and B**). Sera from SUS mice showed significantly higher reactivity against brain lysates than those from CON mice (**Fig. 4B**). Furthermore, the levels of brain-reactive antibodies in the sera correlated with both social avoidance (**Fig. 4C**) and the percentage of immune cell populations controlling antibody production in the cLN (**Figs. S4C and D**). We next stained brain sections from immune-deficient *Rag2^-/-^* mice, which lack endogenous antibodies, with sera from CSDS-exposed mice and detected bound antibodies using anti-mouse IgG secondary antibodies (**Fig. 4D**). Among the samples collected, sera from SUS mice showed the strongest reactivity against brain sections from the NAc (**Fig. 4E**), PFC, and HIP (**Fig. S4E**). Brain sections stained with sera from SUS mice showed a significantly higher fluorescence intensity than those stained with sera from CON mice (**Fig. 4F**). Furthermore, the fluorescence intensity of the brain sections correlated with social avoidance behavior (**Fig. 4G**) and the percentage of GCB and Tfh in the cLN (**Figs. S4F and G**). We also confirmed a correlation of brain reactivity of the sera detected by ELISA and indirect immunofluorescence (**Fig. 4H**). In addition, we measured the reactivity of sera against proteins expressed in the brain via Western blot (**Fig. S4H**). Specific bands were detected when membranes bound to proteins prepared from NAc, PFC, and HIP lysates were incubated with serum from SUS mice. The band patterns differed between lanes with proteins from different brain regions, indicating multiple targets of autoantibodies induced by stress (**Fig. S4I**). To investigate clinical relevance of the findings of brain-reactive antibody induction by stress, we analyzed IgG concentrations and levels of brain-reactive antibodies in clinical samples of sera from healthy controls (HC) and patients with MDD (**Table 1**). Using the Temporal Experience of Pleasure Scale (TEPS), a self-reported measure of pleasure experience (41), we evaluated correlations between brain-reactive antibodies in sera and TEPS anticipatory or consummatory pleasure. Although there was no significant difference of serum IgG concentrations between HC and patients with MDD (**Fig. S5A**), levels of brain-reactive IgG antibodies negatively correlated with the measure of pleasure experience, especially with TEPS anticipatory pleasure (**Figs. 4I, J, and S5B**). These results may provide important evidence indicating clinical relevance of autoimmune responses against the brain induced by stress.

**Fig. 4.**
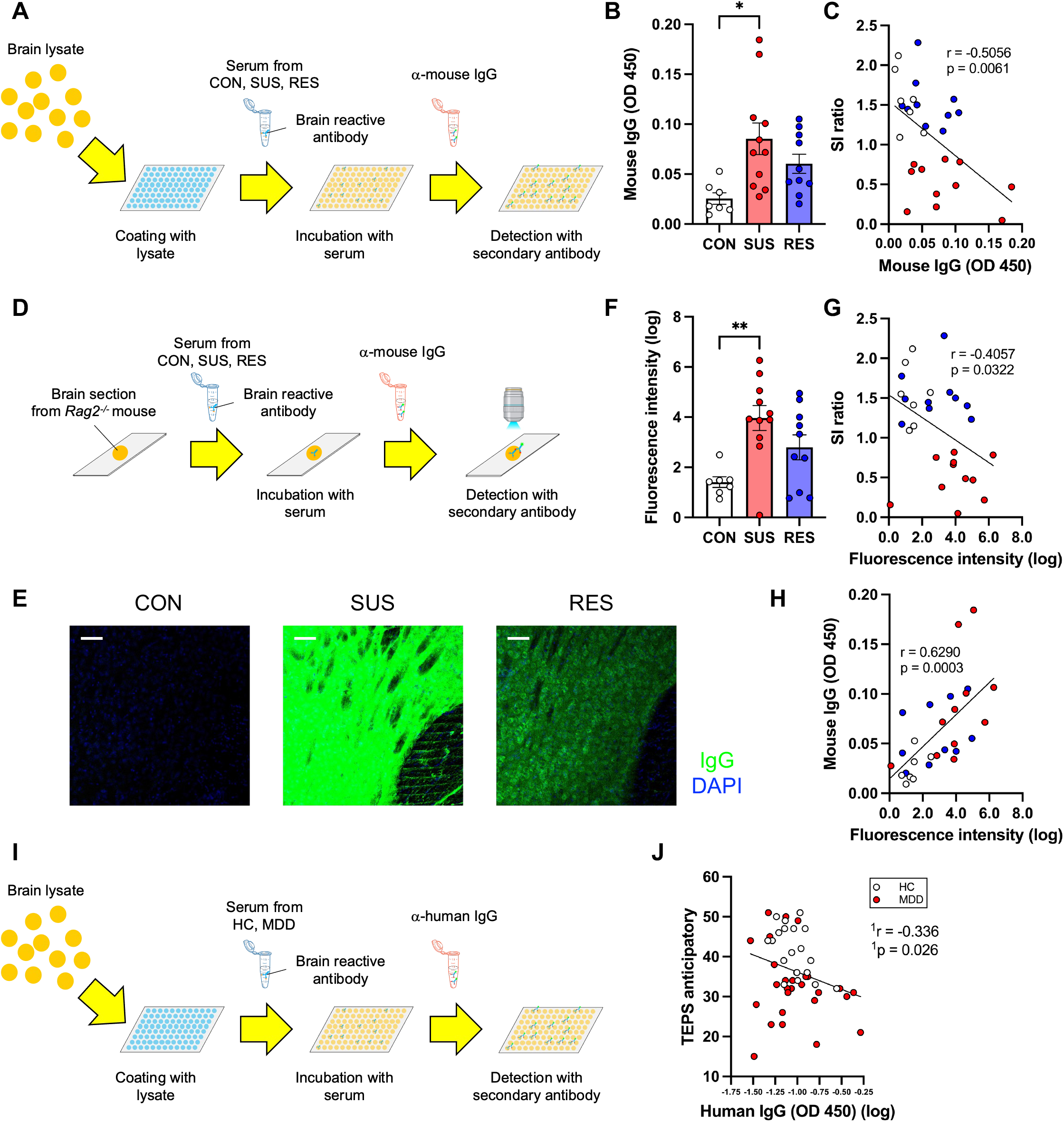
Increase of brain-reactive antibodies in sera from stress-susceptible (SUS) mice and humans with high anhedonia. **A** Experimental outline: Schematic of detection of brain lysate-reactive IgG antibodies in sera from unstressed control (CON), SUS, and stress-resilient (RES) mice by enzyme-linked immunosorbent assay (ELISA). **B** Quantification of brain lysate-reactive IgG antibodies in sera. (CON: n = 7, SUS: n = 11, RES: n = 10). **C** Correlation between brain lysate-reactive IgG antibodies in sera and SI ratio. (CON: n = 7, SUS: n = 11, RES: n = 10). **D** Experimental outline: Schematic of detection of brain section-reactive IgG antibodies in sera by indirect immunofluorescence. **E** Staining of brain sections around the nucleus accumbens (NAc) from immune-deficient *Rag2^-/-^* mice with sera from CON, SUS, and RES mice (green: IgG, blue: DAPI, scale bar: 50 μm). **F** Quantification of fluorescence intensity (CON: n = 7, SUS: n = 11, RES: n = 10). **G** Correlation between fluorescence intensity of brain sections stained with sera and SI ratio (CON: n = 7, SUS: n = 11, RES: n = 10). **H** Correlation between fluorescence intensity of brain sections stained with sera and levels of brain lysate-reactive serum IgG antibodies (CON: n = 7, SUS: n = 11, RES: n = 10). Data represent mean ± standard error of the mean. One-way ANOVA with Bonferroni *posthoc* test (*p < 0.05, **p < 0.01). Correlations were evaluated by Pearson correlation analysis. **I** Experimental outline: Schematic of detection of brain lysate-reactive IgG antibodies in sera from healthy controls (HC) and patients with major depressive disorder (MDD) by ELISA. **J** Correlation between levels of brain lysate-reactive IgG antibodies in sera and the Temporal Experience of Pleasure Scale (TEPS) anticipatory (HC: n = 19, MDD: n = 28). ^1^The partial correlation was calculated to control for the potential confounding variables of age, gender and Body Mass Index (BMI).

### B Cell Depletion before CSDS Promotes Stress Resilience

We further investigated the causal link between antibody responses and stress-susceptibility. Production of autoantibodies is mediated by clonal expansion of autoreactive B cells in the germinal center (42). Thus, we depleted B cells by administering an anti-CD20 antibody before stressing the mice and tested the social behavior of antibody-treated mice after they were subjected to CSDS (**Fig. 5A**). We confirmed the absence of B cells in secondary lymphoid organs using flow cytometry 7 days after anti-CD20 antibody injection in a separate cohort of mice (**Fig. S6A**). Importantly, we observed a marked reduction in the population of B cells in the cLN by anti-CD20 treatment 48 h after the last defeat, indicating that the absence of B cells persisted throughout the experiment (**Figs. 5B and C**). Following CSDS, the B cell-depleted group showed a significantly higher SI ratio than the control IgG-treated group (**Figs. 5D, E and Fig. S6B**), indicating that B cell depletion promoted resilience phenotype. These data suggest that B cells contribute to stress susceptibility in the CSDS model.

**Fig. 5.**
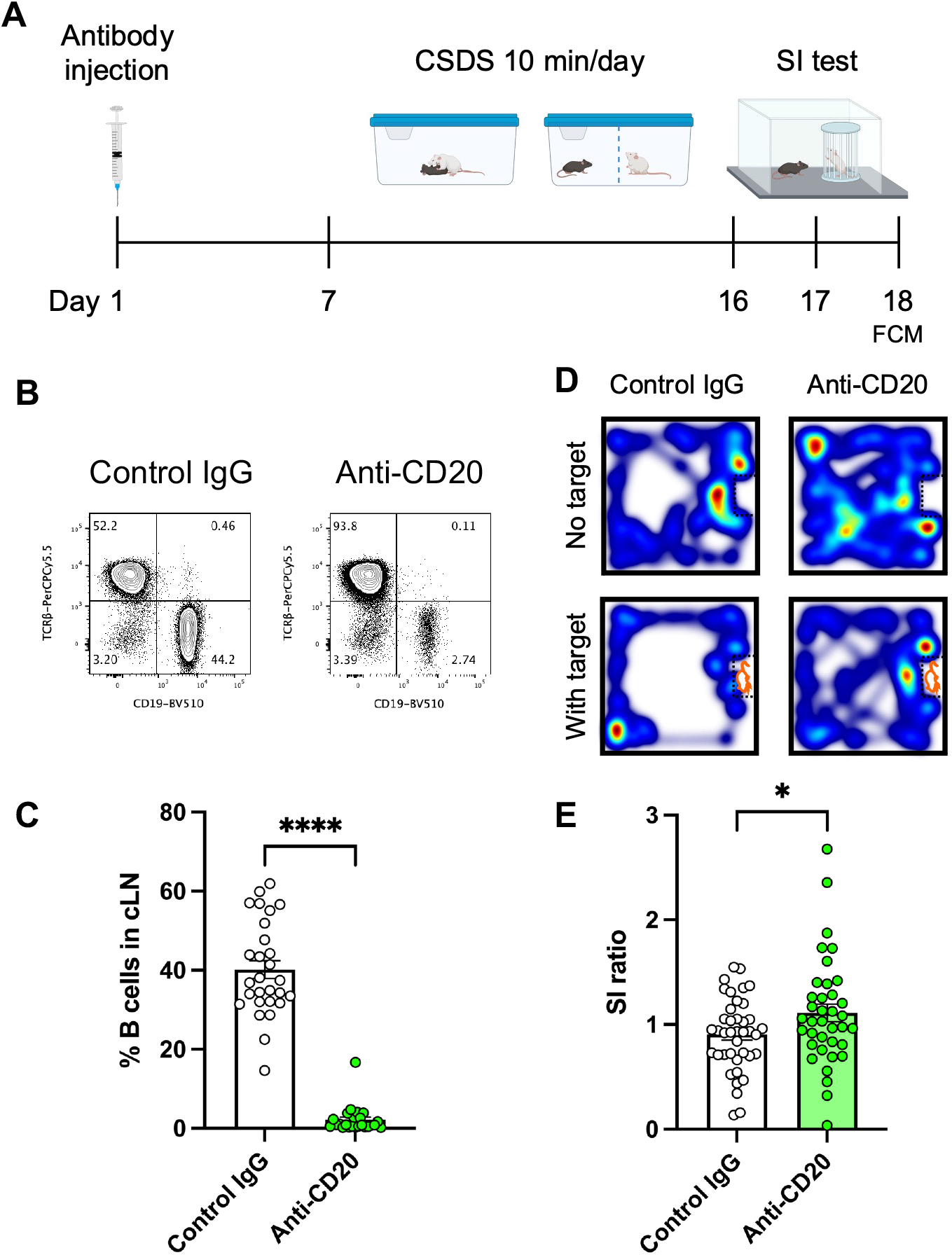
Contribution of B cells to depression-like behavior in the chronic social defeat stress (CSDS) model. **A** Experimental outline: B cell depletion followed by CSDS, social interaction (SI) test and flow cytometry (FCM). **B** Representative flow cytometry plots of lymphocytes from cervical lymph nodes (cLN) at day 18. **C** Analysis of B cell depletion efficiency (Control IgG: n = 28, Anti-CD20: n = 24). **D** Representative heatmaps of mouse behavior during the SI test. **E** Effects of B cell depletion on SI ratio in the CSDS model (Control IgG: n = 40, Anti-CD20: n = 37). Data represent mean ± standard error of the mean. Unpaired t-test (*p < 0.05, ****p < 0.0001).

## Discussion

In this study, we demonstrated mechanisms inducing depression-like behavior mediated by the activation of the adaptive immune system, possibly through the production of autoantibodies targeting antigens in the brain. Previous studies have revealed the critical roles of the innate immune system in depression. Complementing these findings, our current study highlights the important contributions of adaptive immune abnormalities to depression. We observed that social stress induces antibody production in the CSDS model. Specifically, social stress strongly induces germinal center responses in brain-draining lymph nodes and production of brain-reactive antibodies, which correlated with behavioral abnormalities. We conclude that stress susceptibility was regulated by B cells in the CSDS model. These data suggest that CSDS induces autoimmune responses against the brain, which contribute to the pathogenesis of depression-like behavior (**Fig. S7**). Results obtained from this study may provide important evidence for the clinical relevance of stress-induced autoimmune responses against the brain.

Over-activation of immune cell populations such as Tfh, GCB, and PC have been linked to autoimmune diseases (33). cLN have been identified as sites of brain antigen delivery from the CSF through lymphatic vessels in the meninges (43). Recent reports have indicated involvement of cLN in animal models of CNS diseases such as multiple sclerosis, Alzheimer’s disease, and stroke (44–46). To the best of our knowledge, this study is the first to show that social stress induces the activation of antibody responses in the cLN, including strong induction of GCB, Tfh, and PC in cLN from SUS mice. Our results suggest that stress-induced activation of immune cells in these lymph nodes may contribute to depression-like behaviors.

Consistent with a recent study showing cerebrovascular dysfunction in the CSDS model (47), results of the present study revealed that IgG antibodies accumulate around the blood vessels in brain sections from SUS mice. This finding suggests that proteins expressed in the brain vasculature are candidate targets of autoantibodies induced by social stress. Although we reported the production of brain-reactive antibodies in stressed mice and humans with high anhedonia, the exact target antigens of these autoantibodies remain unknown. Identification of the targets will help elucidate the mechanisms by which stress-induced autoantibodies mediate depression.

We found an association between peripheral levels of brain-reactive antibodies and severity of anhedonia in humans. Recent reports have indicated dysfunction of adaptive immune responses, including autoimmunity, in adolescents and young adults with suicidal behavior (48) and activation of B cells in postpartum depression (49); these reports reinforce the clinical relevance of our study findings.

Our results provide novel insights into the mechanisms by which autoimmune responses against the brain mediate depression. Overall, our results suggest that patients with MDD can be classified based on autoantibody production, and offer therapeutic strategies targeting autoimmunity as possible effective treatments.

## Materials and Methods

### Mice

Six to seven week-old male C57BL/6J (Stock#: 000664) mice were purchased from The Jackson Laboratory. Retired breeder male CD-1 mice over 4 months old were purchased from Charles River Laboratories. *Rag2^-/-^* mice were kindly provided by Dr. Miriam Merad. All mice were maintained with *ad libitum* access to food and water. All the experiments were performed in accordance with the National Institutes of Health Guide for Care and approved by the Icahn School of Medicine at Mount Sinai Institutional Animal Care and Use Committee (IACUC).

### Chronic social defeat stress (CSDS)

CSDS was performed as previously described (18). CD-1 mice with aggressive behavior (CD-1 AGG) were selected by inter-male social interactions over 3 consecutive days based on previously described criteria and housed in the social defeat cage (26.7 cm width × 48.3 cm depth × 15.2 cm height, Allentown Inc) 24 h before the start of defeats on one side of a clear perforated Plexiglas divider (0.6 cm × 45.7 cm × 15.2 cm, Nationwide Plastics). Eight week-old experimental C57BL/6J mice were subjected to physical interactions with an unfamiliar CD-1 AGG mouse for 10 min once per day for 10 consecutive days. After antagonistic interactions with the CD-1 AGG mice, experimental mice were removed and housed on the opposite side of the divider, allowing sensory contact over the subsequent 24 h period. Unstressed control (CON) mice were housed two per cage on either side of a perforated divider without being exposed to the CD-1 AGG mice. Experimental mice were singly housed after the last bout of defeat and the social interaction (SI) test was conducted 24 h later.

### Social interaction (SI) test

SI testing was performed as previously described (18). First, mice were placed in a Plexiglas open-field arena with a small wire animal cage placed at one end. Movements were monitored and recorded automatically for 150 s with a tracking system (Ethovision 11.0 Noldus Information Technology) to determine baseline exploratory behavior and locomotion in the absence of a social target (CD-1 AGG mouse). At the end of 150 s, the mouse was removed, and the arena was cleaned with 70 % ethanol. Next, exploratory behavior in the presence of a novel social target inside the small wire animal cage was measured for 150 s and time spent in the interaction zone was analyzed. SI ratio was calculated by dividing the time spent in the interaction zone when the AGG was present by the time spent in the interaction zone when the AGG was absent. All mice with a SI ratio < 1.0 were classified as stress-susceptible (SUS) and all mice with a SI ratio ≥ 1.0 were classified as resilient (RES).

### Flow cytometry

Single cell suspensions were prepared from lymphoid organs (cervical lymph nodes, mesenteric lymph nodes, and spleen) in a staining buffer (0.5% BSA, 2 mM EDTA in PBS). Red blood cells were removed from splenocytes by treatment with 1X lysing buffer (BD Pharm Lyse). After blocking of FcγR (1:200 dilution, 4 □, 30 min), cell surface markers were stained with fluorochrome-labeled antibodies (1:200-400 dilution, 4 □, 30 min). For staining of intracellular markers, cells were fixed and permeabilized using True-Nuclear Transcription Factor Buffer Set (Biolegend) following manufacturers’ instructions. Intracellular markers were stained with fluorochrome-labeled antibodies (1:200-400 dilution, 4 □, 30 min). Dead cells were stained with Fixable Viability Dye eFluor®780 (eBioscience) and excluded from the analyses. Antibodies and other reagents used for the analyses are summarized in Table S1 and S2. Stained cells were acquired on a BD LSRII flow cytometer and obtained data were analyzed with FlowJo software version 10.6.2 (Tree Star).

### B cell depletion

250 μg control IgG antibody (clone: RTK4530, Biolegend) and anti-CD20 antibody (clone: SA271G2, Biolegend) were retro-orbitally injected into seven week-old C57BL/6J mice one week before starting CSDS. Efficiency of B cell depletion was evaluated by analyzing CD19-positive B cells in lymphoid organs by flow cytometry.

### Immunohistochemistry (IHC)

Brains were collected after transcardial perfusion with ice-cold PBS followed by 4% PFA (Electron Microscopy Sciences). Collected brains were post-fixed with 4% PFA (4 □, overnight) and nucleus accumbens (NAc), prefrontal cortex (PFC), and hippocampus (HIP) brain sections were prepared in 50 μm by vibratome (Leica). After blocking with 3% normal Donkey serum with 0.3% Triton X-100 (Sigma) in PBS (RT, 2 h), sections were incubated with primary antibodies (anti-GFAP antibody; 1:1 dilution, anti-CD31 antibody; 1:400 dilution, 4 □, overnight) followed by staining with secondary antibodies (1:300 dilution, RT, 1 h). Nuclei were stained by DAPI (0.5 μg/mL, Molecular Probes). 3D render images were constructed by using IMARIS 9.9 software to create surfaces of each stain based on a threshold applied to all images. Antibodies and other reagents used for the analyses are summarized in Tables S1 and S2. The brain sections were analyzed with a Zeiss LSM 780 confocal microscope.

### Visualization and detection of brain-reactive antibodies

To detect brain-reactive IgG in sera, indirect immunofluorescence analyses were performed. Brain sections (50 μm) from NAc, PFC, and HIP were prepared from *Rag2^-/-^* mice. After blocking as described above, sections were incubated with sera from CON, SUS, and RES mice (1:50 dilution, 4 □, overnight) and bound antibodies were detected and captured with anti-mouse IgG secondary antibodies (1:300 dilution, RT, 1 h) (Jackson ImmunoResearch). The brain sections were imaged and analyzed with a Zeiss LSM 780 confocal microscope. Fluorescence intensity from images were quantified using ImageJ. Briefly, images were first converted to 8-bit greyscale and the region of interest was selected using the drawing/selection tools to create a rectangle, then set as Macro. Macro was then randomly placed to measure fluorescence intensity by calculating the mean value of 3-5 representative fields per sample.

### Brain lysate preparation

Brains were collected after transcardial perfusion with ice-cold PBS. Half brains from mice were mashed in 300 μL of corresponding buffers using the plunger end of a 1 mL syringe. 1X TBS (Fisher) was used as a buffer to prepare lysates for detection of IgG antibodies in the brain lysates. PBS with protease inhibitors (cOmplete™, Mini, EDTA-free Protease Inhibitor Cocktail, Roche) was used as a buffer to prepare lysates for detecting brain lysate-reactive IgG antibodies in sera. Samples were further homogenized by a 1.5 mL pestle. Soluble fractions were collected after centrifugation. Total protein concentrations of the brain lysates were determined using a BCA Protein Assay Kit (Pierce).

### Western blotting

Blood samples were collected from mice and allowed to clot by leaving them undisturbed at room temperature. Sera were collected after centrifugation (2000 x *g*, 4 □, 10 min) and stored at −80 □. Sera were analyzed for the presence of autoantibodies by Western blotting as previously described (50) with some modifications. Briefly, brain lysates were separated by SDS-PAGE and transferred onto membranes (PVDF membrane, Bio-Rad Laboratories). After blocking (Blocking Buffer, ROCKLAND), membranes were incubated with sera (1:500 dilution, 4 □, overnight) and bound antibodies were detected with a IRDye^®^ 800CW-labeled anti-mouse IgG (H+L) secondary antibody (LI-COR) (1:5000 dilution, RT, 40 min). The membranes were analyzed by a LI-COR Odyssey Infrared Imaging system (LI-COR Biosciences).

### Human subjects

Study participants with major depressive disorder (MDD) and healthy controls (HC), as assessed by the Structured Clinical Interview for the Diagnostic and Statistical Manual of Mental Disorders–Fifth Edition (SCID-5) (51) were recruited through the Depression and Anxiety Center for Discovery and Treatment at the Icahn School of Medicine at Mount (ISMMS). The ISMMS review board approved the study, and written informed consent was obtained from all participants prior to any study procedure. Participants were compensated for their time and effort. Subjects provided demographic information and underwent a psychiatric evaluation using the SCID-5 conducted by trained study staff. Participants completed the Quick Inventory of Depressive Symptomatology-SR (QIDS-SR) (52) to measure depressive symptom severity. Anhedonia was assessed using the Temporal Experience of Pleasure Scale (TEPS) (53), a well-validated self-report questionnaire, assessing both, consummatory and anticipatory anhedonia. A higher score on the QIDS-SR indicates higher depression symptom severity and a lower score on the TEPS indicates higher levels of anhedonia. All participants underwent biochemistry and hematological laboratory testing, urine toxicology and pregnancy testing (if applicable). At the time of enrollment, all participants were free of medications known to affect the immune system for at least two weeks. Participants were free of active infections or systemic illness. Subjects with concomitant unstable medical illnesses were excluded. Participants were free of current substances of abuse. On the day of blood draw, patients were fasted for at least 6 h. For serum isolation, blood was drawn into Vacutainer Gold Top 5 mL Silica Gel tubes (BD, #365968). Blood was allowed to clot for 30 min, then centrifuged at 1300 x *g* for 15 min at 4 °C, then aliquoted and stored at −80 °C until further processing.

### Enzyme-linked immunosorbent assay (ELISA)

Total IgG concentrations in sera and brain lysates were analyzed by ELISA following the manufacturers’ instructions. For quantification of brain lysate-reactive IgG antibodies in sera, plates were coated with brain lysates in an ELISA Coating Buffer (Biolegend) instead of mouse or human IgG antibodies.

### Statistical analyses

Details of statistical analyses are described in the figure legends, including type of statistical analysis used, p values, and number of samples. Statistical analyses were performed using GraphPad Prism software (GraphPad Software Inc.) or SPSS (Version 24, IBM Corp., SPSS Inc., Chicago IL, USA). Samples that deviated from the mean by greater than 2 standard deviations were identified as outliers and excluded from the analyses. For the analysis of the data involving human participants, normal distribution of the data was tested using the Kolmogorov-Smirnov Test. If data were not normally distributed, they were log transformed. Partial correlations were calculated to control for the potential confounding variables age, gender and Body Mass Index (BMI). Level of statistical significance was set at p < 0.05.

---

**Fig. S1.**
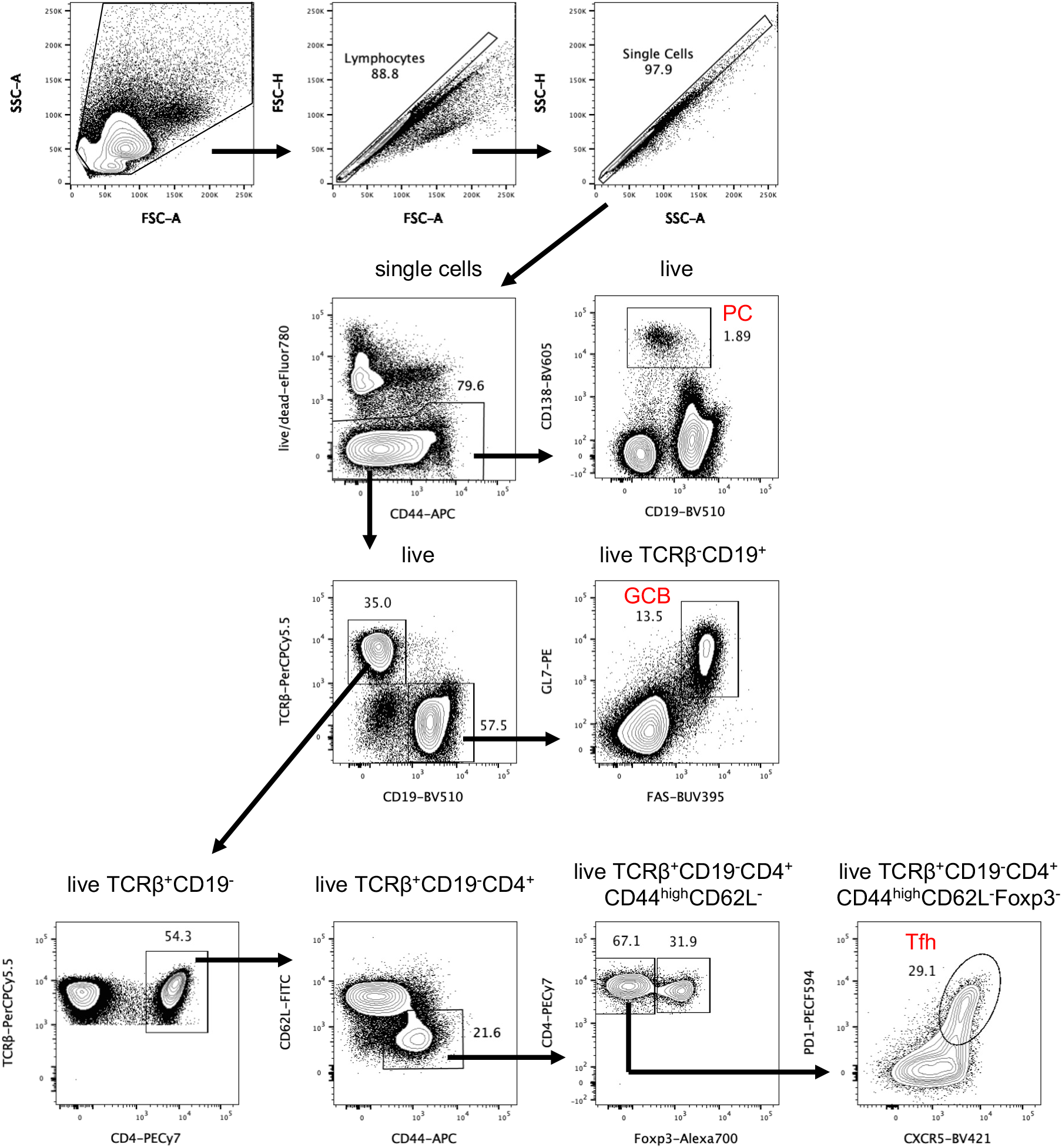
Gating strategy of the flow cytometric analysis of germinal center B cells (GCB), follicular helper T cells (Tfh), and plasma cells (PC). Percentages of immune cell populations were analyzed as described below: GCB: GL7^+^ FAS^+^ in live CD19^+^ TCRβ^-^ cells Tfh: CXCR5^high^ PD1^high^ in live TCRβ^+^ CD19^-^ CD4^+^ CD44^high^ CD62L^-^ Foxp3^-^ cells PC: CD138^+^ in live cells.

**Fig. S2.**
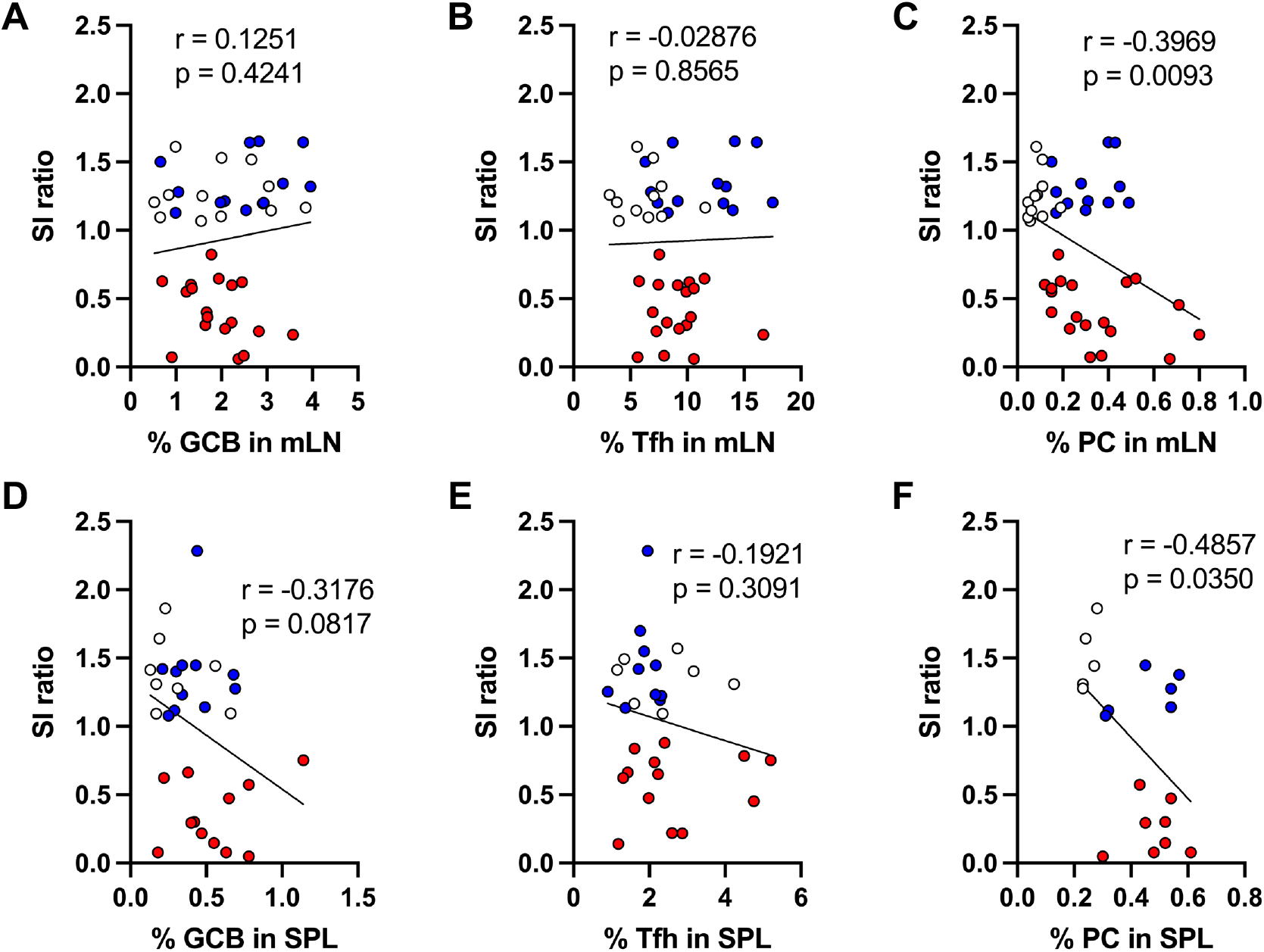
Correlations between social behavior and immune cell populations in mesenteric lymph nodes (mLN) and spleen (SPL) from unstressed control (CON), stress-susceptible (SUS), and stress-resilient (RES) mice. Correlations between social interaction (SI) ratio and percentages of **A** germinal center B cells (GCB) (CON: n = 12, SUS: n = 18, RES: n = 13), **B** follicular helper T cells (Tfh) (CON: n = 11, SUS: n = 18, RES: n = 13), and **C** plasma cells (PC) (CON: n = 11, SUS: n = 19, RES: n = 12) in mLN. Correlations between SI ratio and percentages of **D** GCB (CON: n = 8, SUS: n = 12, RES: n = 11), **E** Tfh (CON: n = 7, SUS: n = 13, RES: n = 10), and **F** PC (CON: n = 5, SUS: n = 8, RES: n = 6) in SPL. Correlations were evaluated by Pearson correlation analysis.

**Fig. S3.**
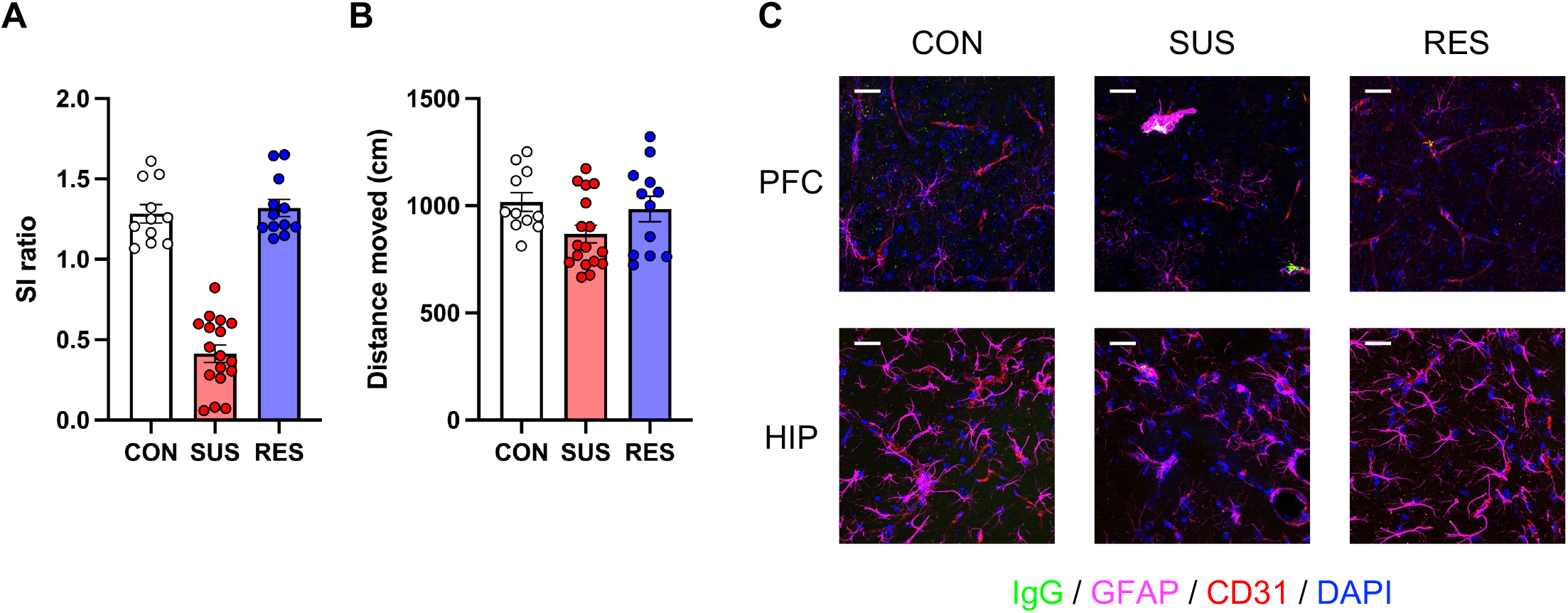
Behaviors and representative brain section images for analyses of IgG antibodies in the brain from unstressed control (CON), stress-susceptible (SUS), and stress-resilient (RES) mice. **A** Social interaction (SI) ratio (CON: n = 11, SUS: n = 17, RES: n = 12). **B** Distance moved during the SI test (No target) (CON: n = 11, SUS: n = 17, RES: n = 12). **C** Staining of IgG antibodies in brain sections of prefrontal cortex (PFC), and hippocampus (HIP) from CON, SUS, and RES mice (green: IgG, magenta: GFAP, red: CD31, blue: DAPI, scale bar: 25 μm). Data represent mean ± standard error of the mean.

**Fig. S4.**
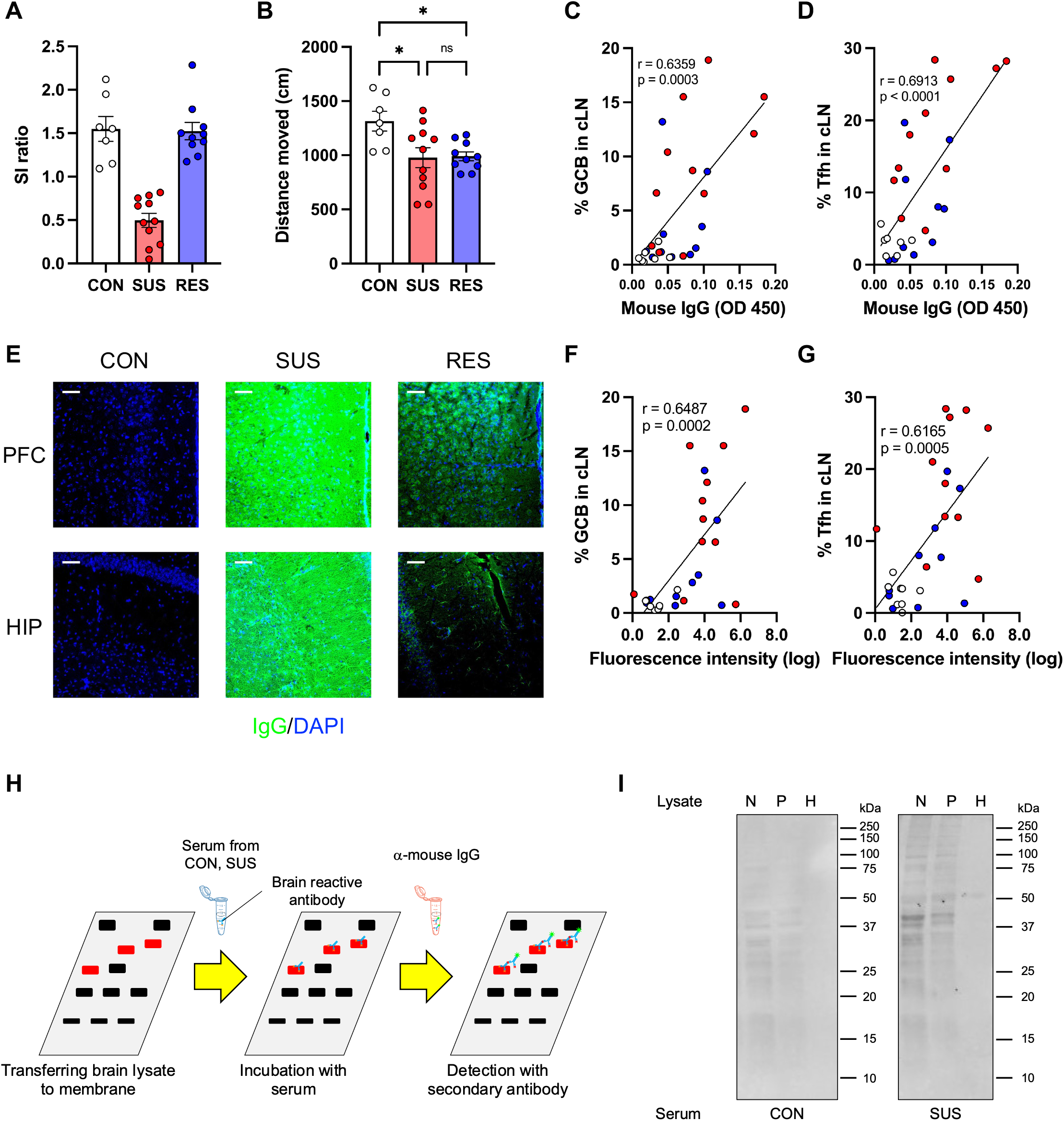
Behaviors, correlations between brain-reactive serum antibody levels and immune cell populations in cervical lymph nodes (cLN), representative brain section images, and Western blotting for analyses of brain-reactive IgG antibodies with sera. **A** Social interaction (SI) ratio of unstressed control (CON), stress-susceptible (SUS), and stress-resilient (RES) mice (CON: n = 7, SUS: n = 11, RES: n = 10). **B** Distance moved during the SI test (No target) (CON: n = 7, SUS: n = 11, RES: n = 10). (C and D) Correlation between levels of brain lysate-reactive serum IgG antibodies and percentages of **C** germinal center B cells (GCB) (CON: n = 7, SUS: n = 11, RES: n = 10), **D** follicular helper T cells (Tfh) (CON: n = 7, SUS: n = 11, RES: n = 10). **E** Staining of brain sections around the prefrontal cortex (PFC) and hippocampus (HIP) from immune-deficient *Rag2^-/-^* mice with sera from CON, SUS, and RES mice (green: IgG, blue: DAPI, scale bar: 50 μm). (F and G) Correlation between fluorescence intensity of brain sections stained with sera and percentages of **F** germinal center B cells (GCB) (CON: n = 7, SUS: n = 11, RES: n = 10), **G** follicular helper T cells (Tfh) (CON: n = 7, SUS: n = 11, RES: n = 10). **H** Schematic for detecting brain-reactive IgG antibodies in sera by Western blotting. **I** Specific bands detected on a membrane with brain lysates from the nucleus accumbens (N), prefrontal cortex (P), and hippocampus (H) incubated with serum from a SUS mouse. Numbers in the figure indicate protein size (kDa). Data represent mean ± standard error of the mean. Oneway ANOVA with Bonferroni *post hoc* test (*p < 0.05, ns: not significant). Correlations were evaluated by Pearson correlation analysis.

**Fig. S5.**
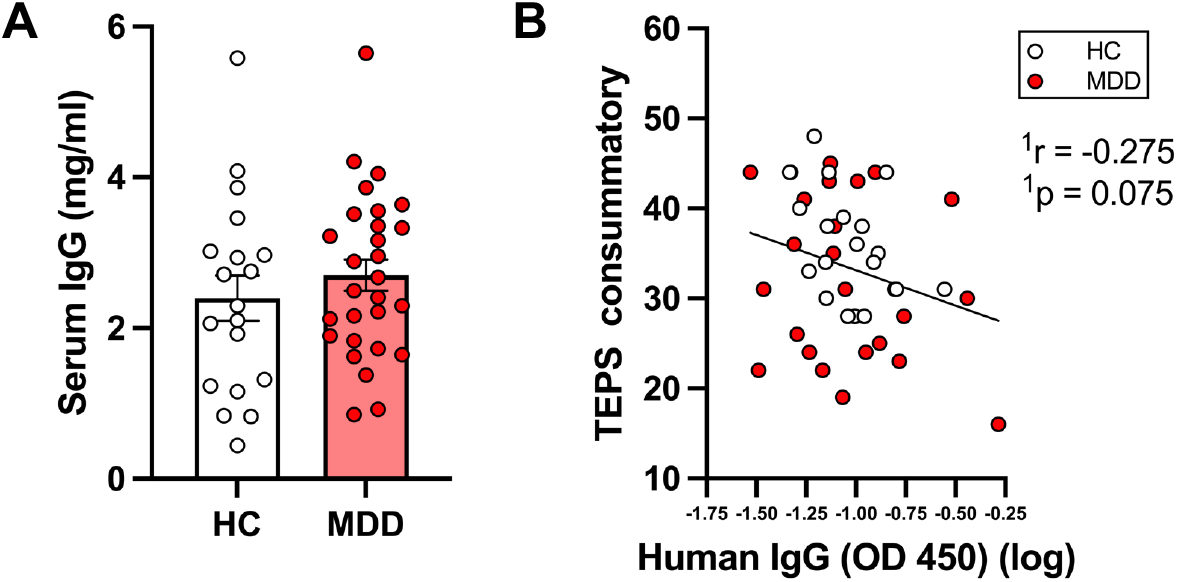
Analyses of serum IgG concentrations and correlation between levels of brain-reactive serum IgG antibodies and anhedonia severity. **A** Total IgG antibody concentrations in sera from healthy controls (HC) and patients with major depressive disorder (MDD). **B** Correlation between levels of brain lysate-reactive IgG antibodies in sera and the Temporal Experience of Pleasure Scale (TEPS) consummatory (HC: n = 19, MDD: n = 27). ^1^The partial correlation was calculated to control for the potential confounding variables of age, gender and BMI (Body Mass Index).

**Fig. S6.**
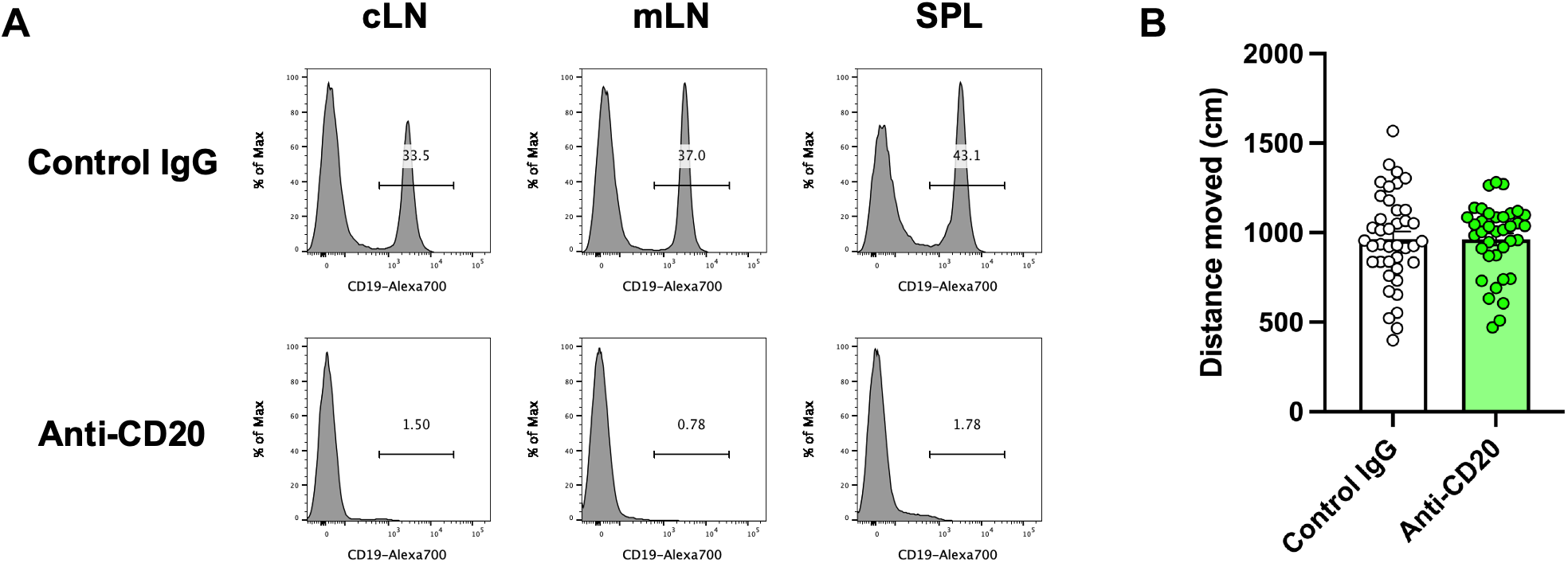
Analyses of B cells in different lymphoid organs and behaviors in B cell depletion experiments. **A** Percentages of B cells in cervical lymph nodes (cLN), mesenteric lymph nodes (mLN), and spleen (SPL) 7 days after control IgG or anti-CD20 antibody treatments. **B** Distance moved during the SI test (No target) (Control IgG: n = 40, Anti-CD20: n = 37). Data represent mean ± standard error of the mean.

**Fig. S7.**
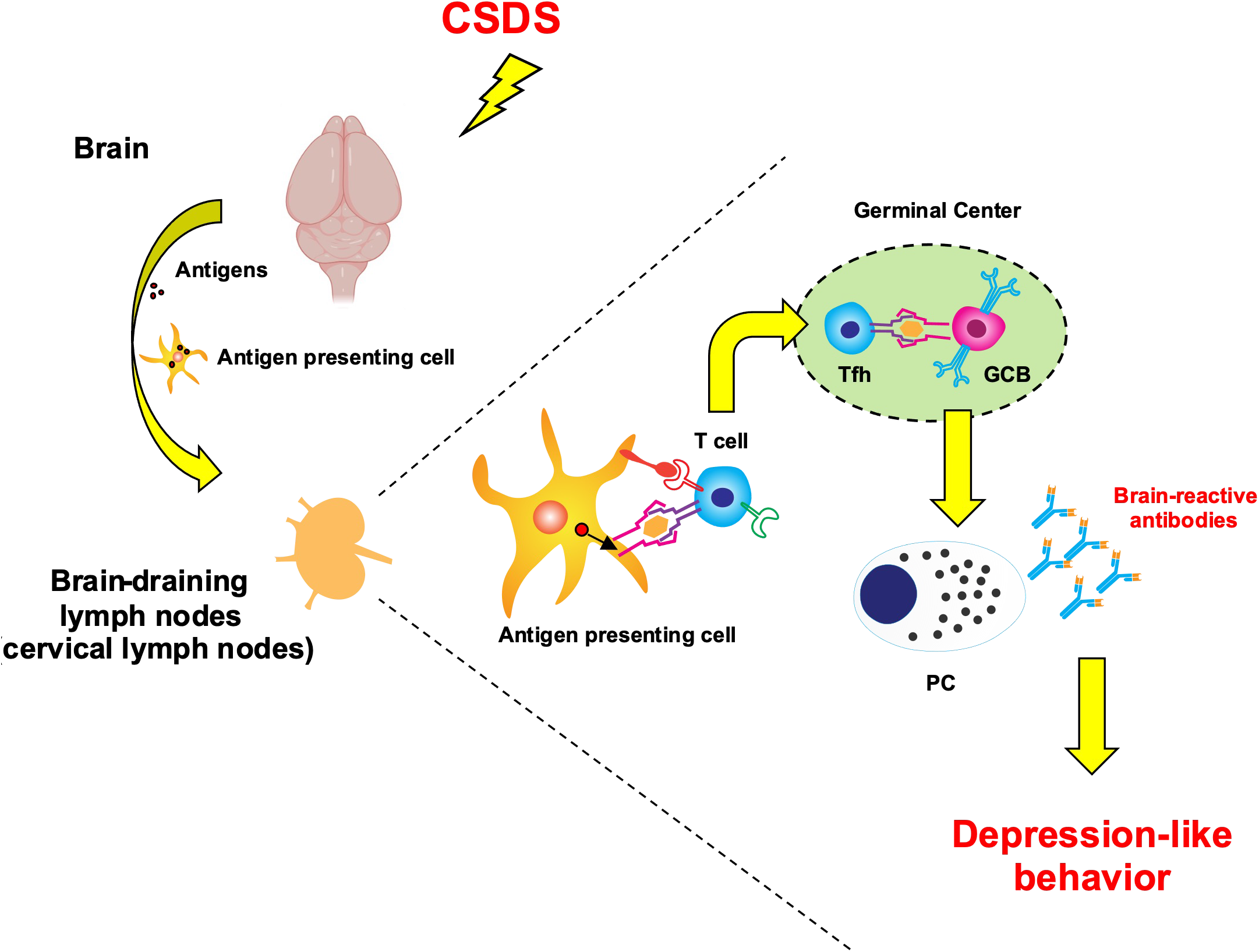
Mechanisms of chronic social defeat stress (CSDS)-induced depression-like behavior mediated by production of autoantibodies against the brain. CSDS induced activation of the germinal center reaction preferentially in brain-draining lymph nodes, leading to production of brain-reactive antibodies. Increase of immune cell populations, such as germinal center B cells (GCB), follicular helper T cells (Tfh), and plasma cells (PC) in cervical lymph nodes was associated with low sociability of stressed mice. Furthermore, elevation of IgG antibody concentrations in the brain and brain-reactive antibody concentrations in sera correlated with the depression-like behavior which was dependent on the presence of B cells.

## References

1. G. S. Malhi, J. J. Mann, Depression. The Lancet 392, 2299–2312 (2018).

2. C. Otte et al., Major depressive disorder. Nat Rev Dis Primers 2, 16065 (2016).

3. A. J. Rush et al., Acute and longer-term outcomes in depressed outpatients requiring one or several treatment steps: a STAR*D report. Am J Psychiatry 163, 1905–1917 (2006).

4. B. N. Gaynes et al., What did STAR*D teach us? Results from a large-scale, practical, clinical trial for patients with depression. Psychiatr Serv 60, 1439–1445 (2009).

5. G. E. Hodes, V. Kana, C. Menard, M. Merad, S. J. Russo, Neuroimmune mechanisms of depression. Nat Neurosci 18, 1386–1393 (2015).

6. A. H. Miller, C. L. Raison, The role of inflammation in depression: from evolutionary imperative to modern treatment target. Nat Rev Immunol 16, 22–34 (2016).

7. R. Dantzer, Cytokine, sickness behavior, and depression. Immunol Allergy Clin North Am 29, 247–264 (2009).

8. J. Barnes, V. Mondelli, C. M. Pariante, Genetic Contributions of Inflammation to Depression. Neuropsychopharmacology 42, 81–98 (2017).

9. W. C. Drevets, G. M. Wittenberg, E. T. Bullmore, H. K. Manji, Immune targets for therapeutic development in depression: towards precision medicine. Nat Rev Drug Discov 10.1038/s41573-021-00368-1 (2022).

10. M. G. Netea, A. Schlitzer, K. Placek, L. A. B. Joosten, J. L. Schultze, Innate and Adaptive Immune Memory: an Evolutionary Continuum in the Host’s Response to Pathogens. Cell Host Microbe 25, 13–26 (2019).

11. C. L. Raison, L. Capuron, A. H. Miller, Cytokines sing the blues: inflammation and the pathogenesis of depression. Trends Immunol 27, 24–31 (2006).

12. G. G. R. Leday et al., Replicable and Coupled Changes in Innate and Adaptive Immune Gene Expression in Two Case-Control Studies of Blood Microarrays in Major Depressive Disorder. Biol Psychiatry 83, 70–80 (2018).

13. N. D. Powell et al., Social stress up-regulates inflammatory gene expression in the leukocyte transcriptome via beta-adrenergic induction of myelopoiesis. Proc Natl Acad Sci U S A 110, 16574–16579 (2013).

14. X. Zheng et al., Chemical dampening of Ly6C(hi) monocytes in the periphery produces anti-depressant effects in mice. Sci Rep 6, 19406 (2016).

15. E. S. Wohleb, N. D. Powell, J. P. Godbout, J. F. Sheridan, Stress-induced recruitment of bone marrow-derived monocytes to the brain promotes anxiety-like behavior. J Neurosci 33, 13820–13833 (2013).

16. A. Niraula, K. G. Witcher, J. F. Sheridan, J. P. Godbout, Interleukin-6 Induced by Social Stress Promotes a Unique Transcriptional Signature in the Monocytes That Facilitate Anxiety. Biol Psychiatry 85, 679–689 (2019).

17. M. L. Pfau et al., Role of Monocyte-Derived MicroRNA106b approximately 25 in Resilience to Social Stress. Biol Psychiatry 86, 474–482 (2019).

18. S. A. Golden, H. E. Covington, 3rd, O. Berton, S. J. Russo, A standardized protocol for repeated social defeat stress in mice. Nat Protoc 6, 1183–1191 (2011).

19. G. E. Hodes et al., Individual differences in the peripheral immune system promote resilience versus susceptibility to social stress. Proc Natl Acad Sci U S A 111, 16136–16141 (2014).

20. C. Menard et al., Social stress induces neurovascular pathology promoting depression. Nat Neurosci 20, 1752–1760 (2017).

21. E. Beurel, L. E. Harrington, R. S. Jope, Inflammatory T helper 17 cells promote depression-like behavior in mice. Biol Psychiatry 73, 622–630 (2013).

22. R. A. Brachman, M. L. Lehmann, D. Maric, M. Herkenham, Lymphocytes from chronically stressed mice confer antidepressant-like effects to naive mice. J Neurosci 35, 1530–1538 (2015).

23. S. M. Clark, J. A. Soroka, C. Song, X. Li, L. H. Tonelli, CD4(+) T cells confer anxiolytic and antidepressant-like effects, but enhance fear memory processes in Rag2(-/-) mice. Stress 19, 303–311 (2016).

24. C. Menard, M. L. Pfau, G. E. Hodes, S. J. Russo, Immune and Neuroendocrine Mechanisms of Stress Vulnerability and Resilience. Neuropsychopharmacology 42, 62–80 (2017).

25. H. Pruss, Autoantibodies in neurological disease. Nat Rev Immunol 21, 798–813 (2021).

26. K. Pape, R. Tamouza, M. Leboyer, F. Zipp, Immunoneuropsychiatry - novel perspectives on brain disorders. Nat Rev Neurol 15, 317–328 (2019).

27. J. Planaguma et al., Human N-methyl D-aspartate receptor antibodies alter memory and behaviour in mice. Brain 138, 94–109 (2015).

28. M. Malviya et al., NMDAR encephalitis: passive transfer from man to mouse by a recombinant antibody. Ann Clin Transl Neurol 4, 768–783 (2017).

29. J. Dalmau et al., An update on anti-NMDA receptor encephalitis for neurologists and psychiatrists: mechanisms and models. The Lancet Neurology 18, 1045–1057 (2019).

30. C. Kowal et al., Human lupus autoantibodies against NMDA receptors mediate cognitive impairment. Proc Natl Acad Sci U S A 103, 19854–19859 (2006).

31. R. J. Ludwig et al., Mechanisms of Autoantibody-Induced Pathology. Front Immunol 8, 603 (2017).

32. D. S. W. Lee, O. L. Rojas, J. L. Gommerman, B cell depletion therapies in autoimmune disease: advances and mechanistic insights. Nat Rev Drug Discov 20, 179–199 (2021).

33. S. G. Tangye, C. S. Ma, R. Brink, E. K. Deenick, The good, the bad and the ugly - TFH cells in human health and disease. Nat Rev Immunol 13, 412–426 (2013).

34. M. Valles-Colomer et al., The neuroactive potential of the human gut microbiota in quality of life and depression. Nat Microbiol 4, 623–632 (2019).

35. D. Li et al., 3beta-Hydroxysteroid dehydrogenase expressed by gut microbes degrades testosterone and is linked to depression in males. Cell Host Microbe 10.1016/j.chom.2022.01.001 (2022).

36. D. B. McKim et al., Sympathetic Release of Splenic Monocytes Promotes Recurring Anxiety Following Repeated Social Defeat. Biol Psychiatry 79, 803–813 (2016).

37. W. P. Lafuse et al., Exposure to a Social Stressor Induces Translocation of Commensal Lactobacilli to the Spleen and Priming of the Innate Immune System. J Immunol 198, 2383–2393 (2017).

38. A. Louveau et al., Structural and functional features of central nervous system lymphatic vessels. Nature 523, 337–341 (2015).

39. A. Aspelund et al., A dural lymphatic vascular system that drains brain interstitial fluid and macromolecules. J Exp Med 212, 991–999 (2015).

40. M. D. Sweeney, Z. Zhao, A. Montagne, A. R. Nelson, B. V. Zlokovic, Blood-Brain Barrier: From Physiology to Disease and Back. Physiol Rev 99, 21–78 (2019).

41. Z. Li et al., The structural invariance of the Temporal Experience of Pleasure Scale across time and culture. Psych J 7, 59–67 (2018).

42. J. Suurmond, B. Diamond, Autoantibodies in systemic autoimmune diseases: specificity and pathogenicity. J Clin Invest 125, 2194–2202 (2015).

43. Z. Papadopoulos, J. Herz, J. Kipnis, Meningeal Lymphatics: From Anatomy to Central Nervous System Immune Surveillance. J Immunol 204, 286–293 (2020).

44. A. Louveau et al., CNS lymphatic drainage and neuroinflammation are regulated by meningeal lymphatic vasculature. Nat Neurosci 21, 1380–1391 (2018).

45. S. Da Mesquita et al., Functional aspects of meningeal lymphatics in ageing and Alzheimer’s disease. Nature 560, 185–191 (2018).

46. E. Esposito et al., Brain-to-cervical lymph node signaling after stroke. Nat Commun 10, 5306 (2019).

47. M. L. Lehmann, C. N. Poffenberger, A. G. Elkahloun, M. Herkenham, Analysis of cerebrovascular dysfunction caused by chronic social defeat in mice. Brain Behav Immun 88, 735–747 (2020).

48. M. K. Jha et al., Dysfunctional adaptive immune response in adolescents and young adults with suicide behavior. Psychoneuroendocrinology 111, 104487 (2020).

49. J. Guintivano et al., Transcriptome-wide association study for postpartum depression implicates altered B-cell activation and insulin resistance. Mol Psychiatry 10.1038/s41380-022-01525-7 (2022).

50. W. Jiang, M. S. Anderson, R. Bronson, D. Mathis, C. Benoist, Modifier loci condition autoimmunity provoked by Aire deficiency. J Exp Med 202, 805–815 (2005).

51. M. B. First, J. B. W. Williams, R. S. Karg, R. L. Spitzer, Structured Clinical Interview for DSM-5 Research Version SCID-5-RV (American Psychiatric Association, Arlington, VA, 2015).

52. A. J. Rush et al., The 16-Item Quick Inventory of Depressive Symptomatology (QIDS), clinician rating (QIDS-C), and self-report (QIDS-SR): a psychometric evaluation in patients with chronic major depression. Biol Psychiatry 54, 573–583 (2003).

53. D. E. Gard, A. M. Kring, M. G. Gard, W. P. Horan, M. F. Green, Anhedonia in schizophrenia: distinctions between anticipatory and consummatory pleasure. Schizophr Res 93, 253–260 (2007).

