## Supplementary material for "Social stress induces autoimmune responses against the brain to promote stress susceptibility": Table 1

|  | **HC (n = 19)** | **MDD (n = 28)** | **Statistics** |
| --- | --- | --- | --- |
| **Age** | 39.95 ± 9.65 | 33.43 ± 9.71 | t(45) = -2.264,  *p* = 0.028 |
| **Gender (m/f)** | (10/9) | (11/17) | χ2 = 0.816,  *p* = 0.390 |
| **BMI** | 25.72 ± 3.79 | 24.57 ± 5.48 | t(45) = -0.789,  *p* = 0.434 |
| **QIDS-SR total** | 1.63 + 1.71 | 13.54 + 4.80 | t(45) = 10.353,  *p* < 0.0001 |

**Table 1. Sociodemographic variables and clinical data.** Statistics: two-tailed Student’s t-test (for age, BMI and QIDS-SR), Pearson's chi-squared test (Gender). Abbreviations: BMI: Body Mass Index; f: Female; HC: Healthy controls; m: Male; MDD: Major depressive disorder; QIDS-SR: Quick Inventory of Depressive Symptomatology-SR.
